# Assessing heterogeneity in spatial data using the HTA index with applications to spatial transcriptomics and imaging

**DOI:** 10.1101/2021.02.28.433250

**Authors:** Alona Levy-Jurgenson, Xavier Tekpli, Zohar Yakhini

## Abstract

**Motivation:** Tumour heterogeneity is being increasingly recognised as an important characteristic of cancer and as a determinant of prognosis and treatment outcome. Emerging spatial transcriptomics data hold the potential to further our understanding of tumour heterogeneity and its implications. However, existing statistical tools are not sufficiently powerful to capture heterogeneity in the complex setting of spatial molecular biology.

**Results:** We provide a statistical solution, the HeTerogeneity Average index (HTA), specifically designed to handle the multivariate nature of spatial transcriptomics. We prove that HTA has an approximately normal distribution, therefore lending itself to efficient statistical assessment and inference. We first demonstrate that HTA accurately reflects the level of heterogeneity in simulated data. We then use HTA to analyse heterogeneity in two cancer spatial transcriptomics datasets: spatial RNA sequencing by 10x Genomics and spatial transcriptomics inferred from H&E. Finally, we demonstrate that HTA also applies to 3D spatial data using brain MRI. In spatial RNA sequencing we use a known combination of molecular traits to assert that HTA aligns with the expected outcome for this combination. We also show that HTA captures immune-cell infiltration at multiple resolutions. In digital pathology we show how HTA can be used in survival analysis and demonstrate that high levels of heterogeneity may be linked to poor survival. In brain MRI we show that HTA differentiates between normal ageing, Alzheimer’s disease and two tumours. HTA also extends beyond molecular biology and medical imaging, and can be applied to many domains, including GIS.

**Availability:** <attached; publicly available upon acceptance>.

**Supplementary information:** Supplementary data are available at *Bioinformatics* online.

## 1 Introduction

This study provides a novel solution for characterising and statistically assessing spatial heterogeneity. Recently, there has been growing evidence that phenotypical and clonal heterogeneity may play a crucial role in tumour biology and in affecting cancer progression and treatment outcome [12, 2]. Cancer cells differ in molecular characteristics such as mutations, gene expression and copy number aberrations. These differences, which define the concept of clonality in tumours, are a potentially detrimental hallmark of cancer. In particular, tumour sub-populations may possess a unique combination of molecular traits that enables them to evade treatment [6]. The heterogeneous environment arising from such sub-populations has been mainly investigated through bulk measurements. However, bulk measurements lack the spatial dimension, which may harbour potentially critical information. For example, the evolutionary dynamics of cancer may result in tumour subclones residing in distinct microhabitats that support the development of therapy-resistant populations [7]. In glioblastoma, differences in copy number alterations and somatic mutations were observed when assessing different tumour microenvironments: EGFR-amplified cancer cells were mainly found in poorly vascularised regions, whereas PDGFRA-amplified cancer cells were observed in close proximity to endothelial cells [11]. The spatial distribution of immune-cells among tumour cells has a long-standing role in diagnosis [8], and was proven useful in predicting prognosis and treatment response in multiple cancer types and molecular settings [21, 15]. Recent advances in spatial transcriptomics, including technology developed for direct measurement (e.g., Visium spatial RNA-sequencing (RNA-seq) by 10x Genomics [1]), as well as approaches for inferring such information from digital pathology images [5, 10], have accentuated the interest in analysing molecular heterogeneity from a spatial perspective [3, 10, 14], with some studies already indicating its potential clinical utility [14, 10].

To support such analyses, we have developed a statistical tool that measures the level of spatial heterogeneity – the HeTerogeneity Average index (HTA). We demonstrate its use using synthetic data, two spatial transcriptomics datasets and four brain MRI scans. We also demonstrate its applicability to other domains.

Several methods have been recently adopted from other fields, mainly ecology, to assist in the quantitative analysis of the spatial heterogeneity of molecular measurements [20]. However, such methods, originating from other fields, do not easily extend to complex biological environments. First, they were mostly designed for univariate and bivariate analyses. For example, Morisita-Horn [17] is a measure of overlap between two types of elements, such as two species. It has been used in [13] to measure the colocalisation of immune and cancer cells in breast cancer; Moran’s I, and the more recent *q*-statistic [19], both originating in ecology, measure the spatial auto-correlation and spatial stratified heterogeneity (respectively) of a single attribute with respect to neighbouring locations in space. Another method, Ripley’s K [18], determines whether a single attribute is dispersed, clustered, or randomly distributed in the target spatial environment. Since we are interested in analysing complex biological environments, with many molecular traits, univariate and bivariate methods fall short of providing an adequate solution (as demonstrated in Section 4). Moreover, these methods may also be difficult to interpret or complex to use (e.g., including edge-correction and radius parameters as in Ripley’s K). Importantly, little is known about the distribution of the null hypothesis for the vast majority of these methods. For Morisita-Horn and Ripley’s K, for example, *p*-values are empirically estimated using Monte-Carlo simulations, which are computationally expensive and less accurate compared to methods based on a known null distribution.

The method we propose in this paper, HTA, which is based on Shannon’s entropy, addresses these shortcomings. First, HTA is multivariate, allowing it to capture a richer representation of heterogeneity, even in the bivariate case (see Section 4, Figure 10); second, it lends itself to easier interpretation since it is based on the notion of entropy; and third, for a fixed set of traits, it requires only a single input parameter. Importantly, the HTA distribution, under a null model, can be well characterised and it thus facilitates efficient statistical assessment and inference.

**Fig. 10.**
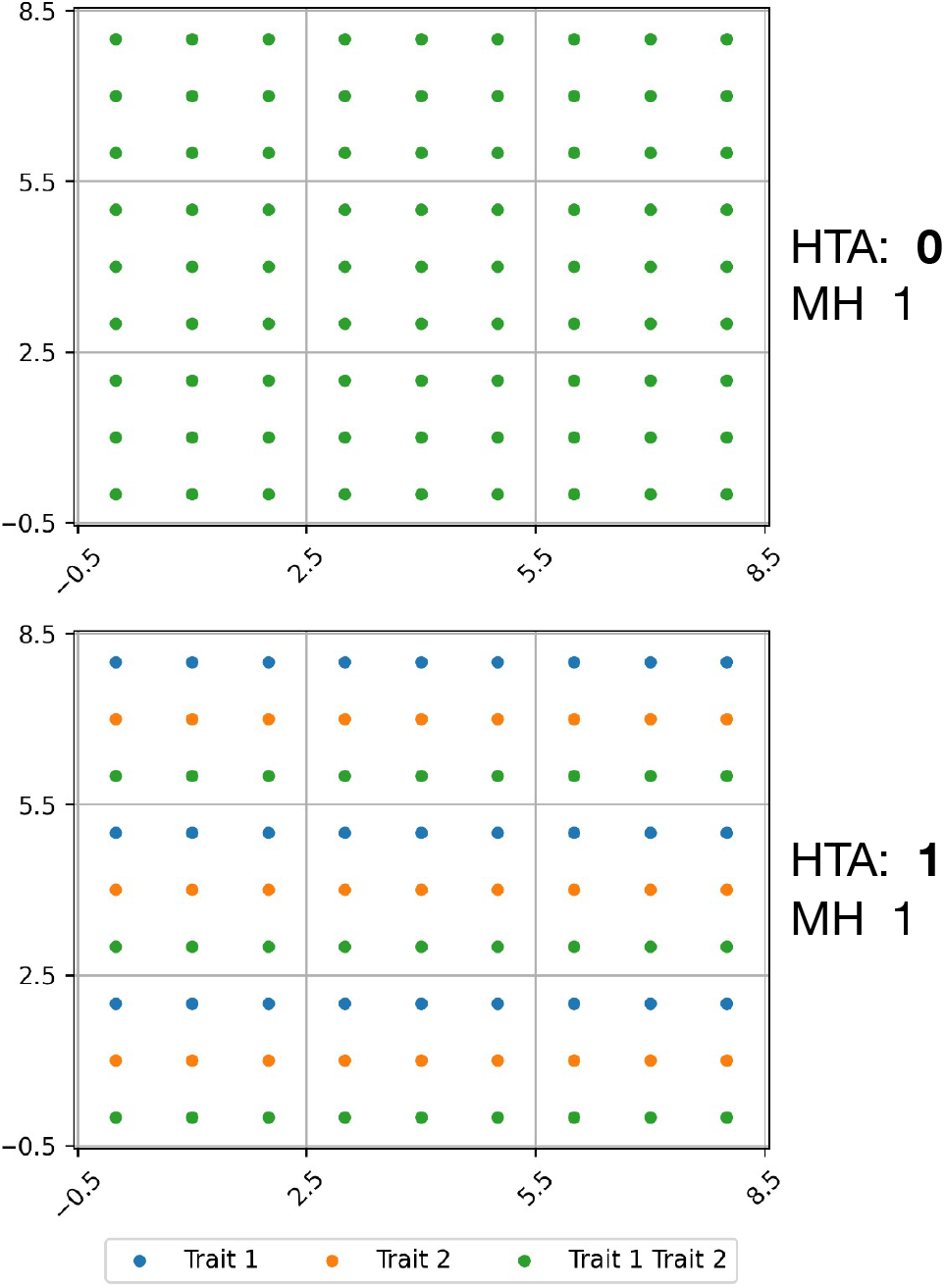
HTA captures a richer representation of heterogeneity, allowing it to differentiate between types of overlap (a proxy used for heterogeneity), even in the bivariate case.

## 2 Methods

In this section we introduce HTA - the HeTerogeneity Average index. We first (Section 2.1) define an index called HTI (HeTerogeneity Index – a variation of Shannon’s entropy) which we will use to measure heterogeneity at a local level. The HTA index (Section 2.2), representing heterogeneity at the whole sample level, will be based on averaging local HTIs. Finally (Section 2.3), we prove that HTA has an approximately normal distribution.

### 2.1 HTI

We first define a heterogeneity index, HTI [10], on which HTA is based. HTI is a variation of Shannon’s entropy. Formally:

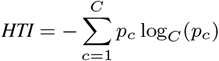

where *C* is the number of non-empty trait combinations that may be observed in the analysed sample, which typically equals 2^|*traits*|^ − 1 (the number of subsets excluding the empty set); and *p*_*c*_ is the proportion of spatial positions for which exactly all traits in combination *c* manifest. A spatial position, for instance, could be a barcoded spot in spatial transcriptomics data or a tile derived from a pathology whole-slide image as in [5, 10].

For example, for two traits, e.g. FOXA1 and MKI67, whose gene expression levels were spatially resolved to a whole slide image from a breast cancer sample, we have *C* = 3 for 3 possible non-empty trait combinations: FOXA1 (only), MKI67 (only) and Both. If the tissue is homogeneous with nearly all of its sections falling into one of these three options (say Both), then *p*(*Both*) *≅* 1, *p*(*FOXA*1) *≅* 0, *p*(*MKI*67) *≅* 0, and HTI is 0. If, however the tissue is heterogeneous with 1/3 of the tiles falling into each option then: *p*(*Both*) = 1*/*3, *p*(*FOXA*1) = 1*/*3, *p*(*MKI*67) = 1*/*3 and HTI is 1. The logarithm base *C* guarantees that HTI falls within [0, 1]. In this case, a high HTI indicates there may be two or more phenotypically different cell types, whereas a low HTI would reflect single phenotypical dominance.

While HTI was shown to capture heterogeneity at a global level [10], it is agnostic to the within-tissue distribution of the trait combinations. For example, the sample in Figure 2 (left), and the sample in Figure 2 (right) have different spatial distributions of the same elements, but HTI is 1 in both cases. This is expected since HTI is a global measure of heterogeneity. However, there is clearly a difference in heterogeneity at the local level, which may have important clinical implications. Our HTA index, described below, which uses HTI at the local level, is designed to capture this difference. Indeed, as noted in Figure 2, the corresponding HTA scores are 0 (homogeneous) on the left and 1 (heterogeneous) on the right.

**Fig. 2.**
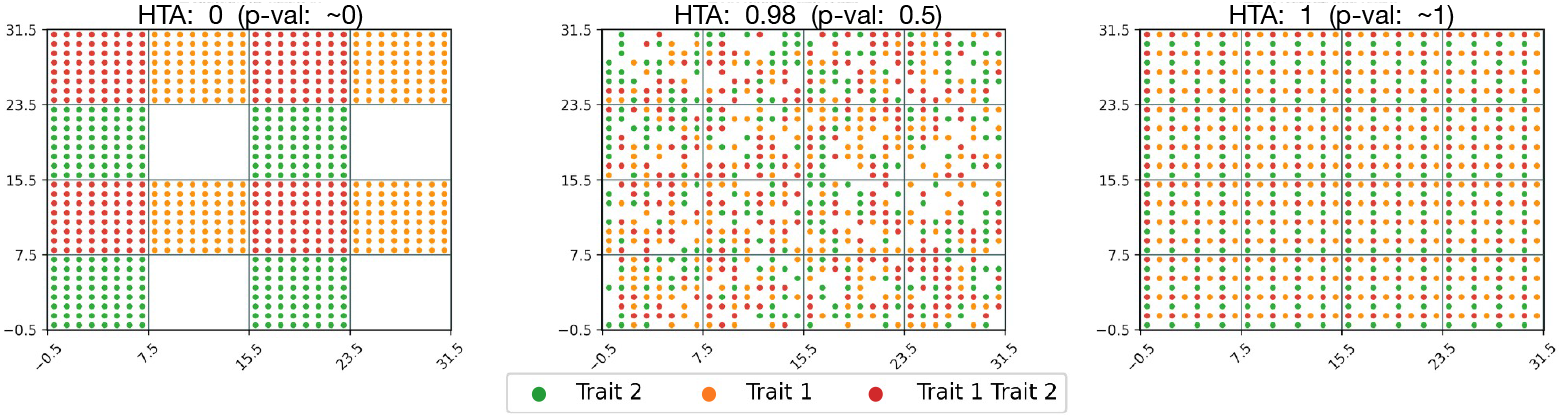
HTA applied to synthetic data across three different distributions (homogeneous, random heterogeneous, deterministic heterogeneous from left-to-right). Region size and trait proportions are held constant (8 and 0.5 resp.). HTA *p*-values: (left) significant homogeneity (*p ≅* 0); (middle) not significant (*p* = 0.23); (right) significant heterogeneity (1-*p ≅* 0).

### 2.2 HTA

#### 2.2.1 HTA definition

HTA, HeTerogeneity Average index, is essentially a weighted average of HTIs across a defined set of spatial regions of a sample (Figure 1 depicts such regions). To formally describe HTA, we first define what regions of a matrix are, and then use these to define HTA.

**Fig. 1.**
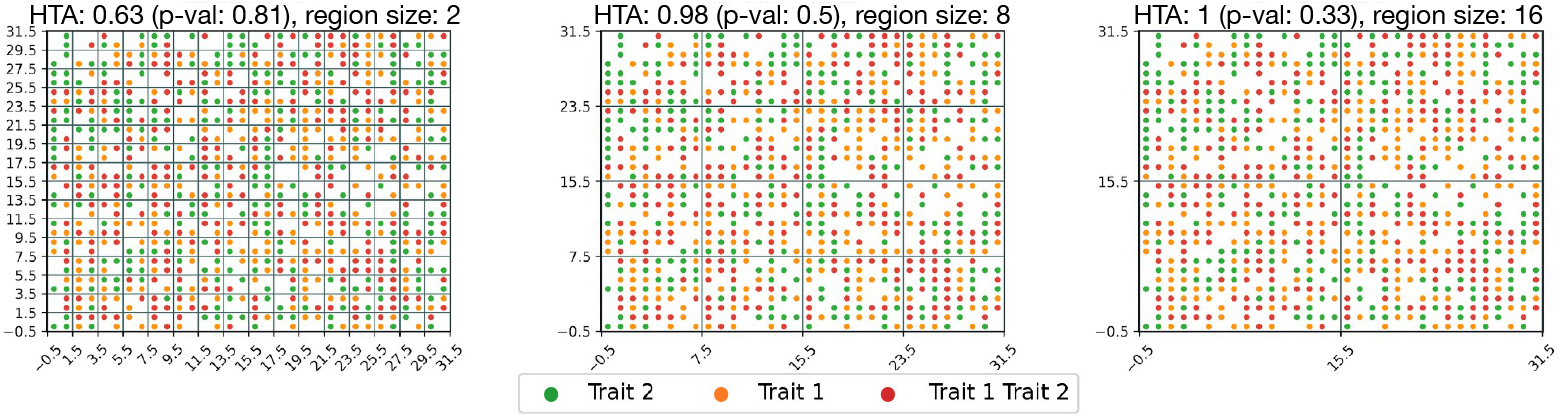
HTA applied to synthetic random data of shape (32, 32) across three different region sizes (2,8,16 left-to-right). The data (dot location and color) is held constant across all three. *p*-values demonstrate heterogeneity since *H*_0_ is not rejected (HTA *p*-value > 0.3) for all region sizes.

Consider a matrix *M*, where each entry corresponds to a spatial location in the sample (e.g. a barcoded spot from spatial RNA-seq data or a single tile from a pathology image [10]) and indicates which of the *C* trait combinations is present therein (or ‘None’ otherwise). We will call such a matrix a trait-combination matrix.

##### HTA

Let *M* |_*G*_ = {*M*_1_, *M*_2_, …, *M*_*R*_} be the set of regions obtained by applying grid *G* to some trait-combination matrix *M*. Let {*n*_1_, *n*_2_, …, *n*_*R*_} be the corresponding number of entries in each region that are not ‘None’. Then we define:

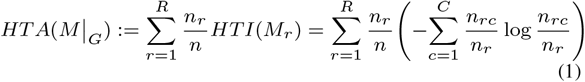

where *n*_*rc*_ is the number of entries in region *r* that manifest trait combination *c* ∈ *{*1, …, *C}*, and *n* is the total number of entries, in the entire matrix, that manifest at least one trait.

Note that since *n*_*r*_ is the number of entries in region *r* that are not ‘None’, this means that 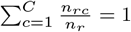, for all *r*. Empty regions (where all elements in the region are ‘None’) are discarded.

For example, in spatial RNA-seq, each entry in *M* that is included in *n* represents a barcoded spot that manifests at least one trait (e.g., the coloured dots in Figure 3D), and each included region of *M* contains several such barcoded spots (e.g., the non-empty regions bordered by the grid lines in the same figure). In digital pathology, where each whole-slide image is divided into thousands of smaller images (tiles), each entry represents a single tile in which at least one trait is present and each region of *M* contains several such tiles.

**Fig. 3.**
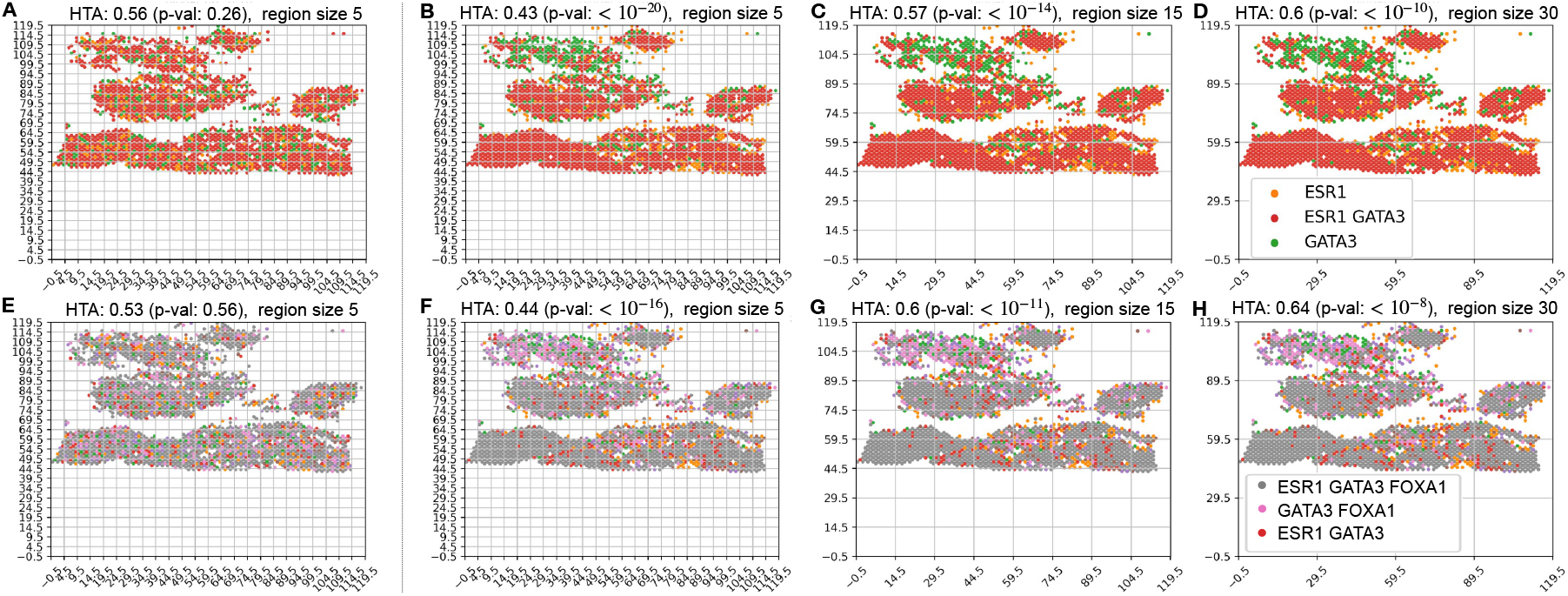
Heterogeneity maps and corresponding HTAs for: (B-D) two traits: ESR1 and GATA3; (F-H) three traits: ESR1, GATA3 and FOXA1. Each color represents the manifestation of a different trait combination. In B-D, red means that ESR1 and GATA3 are highly expressed (above their respective medians), green – only GATA3 is highly expressed and orange – only ESR1 is highly expressed. In F-H, due to the large number of trait-combinations, we note here the most common trait combinations and provide the full legend in Supplementary S4: grey – all three traits are highly expressed, pink – GATA3 and FOXA1 are highly expressed, red – ESR1 and GATA3 are highly expressed. HTA is significantly homogeneous at all region sizes (5, 15 and 30) in both settings (HTA *p*-values < 10^−8^). This aligns with the expected outcome for this cancer type (Luminal B breast cancer, for which the cancer cells highly express these three transcription factors); (A) and (E): the resulting heterogeneity maps if the respective trait-combinations were randomly distributed (*H*_0_). HTA *p*-values are 0.26 and 0.56 (A, E respectively) at region size 5, as expected under *H*0.

We note that HTA monotonically decreases with grid refinement. This is similar to the fact that:

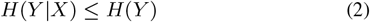

for any random variables *X* and *Y* (for a proof see Supplementary S1). Indeed, we observe this in Figure 1, where HTA decreases from right-to-left. We note that since region-based methods are inherently sensitive to region size, HTA’s monotonicity provides an added advantage since it guarantees an ordering one can expect to observe when moving between region sizes. In Figure 4, for example, we can see that for the largest region size (D), the null hypothesis of heterogeneity is not rejected (HTA *p* = 0.1), whereas at the smaller region sizes (B-C) it is. Knowing that HTA monotonically decreases with grid refinement, a user may be inclined to test finer grids before concluding that the sample is heterogeneous with respect to the mutual spatial distribution of T-cells among HER2 cells.

**Fig. 4.**
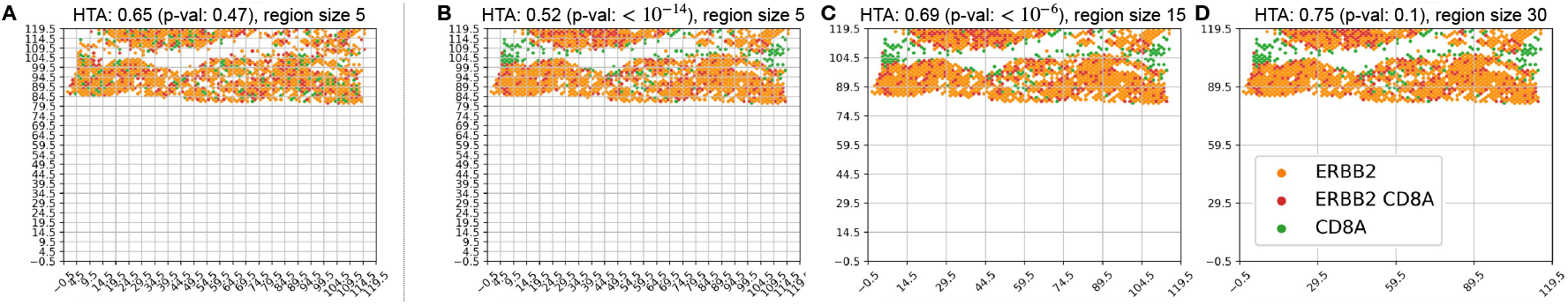
(B-D) Heterogeneity maps and HTA depicting the mutual distribution of ERBB2 (HER2) and CD8A (T-cells) for the bottom half of Figure 3, chosen due to indications of high tumour content in that portion (see Section 3.2). (A) The resulting heterogeneity map if ERBB2 and CD8A were randomly distributed (*H*_0_). HTA is: (B-C) significantly homogeneous *p* < 10^−6^; (D) heterogeneous *p* = 0.1 (*H*_0_ not rejected); (A) heterogeneous *p* = 0.47.

### 2.3 HTA *p*-value

We compute the HTA *p*-value under the null model in which all trait combinations are uniformly distributed across the tissue sample, as in Figures 3 A, E and 4 A (i.e., a random permutation of the exact trait combinations present within the tissue sample, naturally preserving the observed frequencies).

#### 2.3.1 Equal-weight regions

If we assume that all regions contain the same number of entries, we obtain that HTA is normally distributed, by the classical central-limit theorem (Lindeberg–Lévy CLT). Formally, we denote *X*_*r*_, *r* = 1, …, *R* to be *HT I*(*M*_*r*_). Then, under the null hypothesis, the random variables *X*_*r*_ are iid. Therefore, by the CLT, their mean (HTA) is normally distributed:

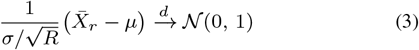

where *µ* and *σ* are the mean and standard-deviation of *X*_*r*_, under the null model.

In our case this means:

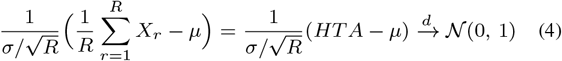

*µ* and *σ* depend on both the region size and the distribution of trait combinations in the matrix *M*. We can estimate these quantities from simulations of *M* under the null model. For a limited region size, we can also compute these quantities precisely by running an exhaustive search across the permutations of trait combinations in a single region to obtain all possible values for *X*_*r*_ and its resulting distribution.

Using this approach for two traits (3 non-empty trait combinations, *C* = 3) and region sizes *s* = 2 and *s* = 3 (yielding square regions with 2^2^ = 4 and 3^3^ = 9 elements, respectively) we obtain, for example, that for sufficiently large values of *R* the following holds:

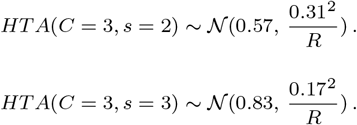

#### 2.3.2 Weighted regions

For actual data, the number of entries in each region may vary as a function of the positions in which entries of *M* are empty. Their corresponding HTIs are therefore no longer identically distributed under the null hypothesis. Specifically, we have different means and stds for the HTI of each of the regions, indexed by *r*, which we denote by *µ*_*r*_ and *σ*_*r*_, respectively. For example, a region with only one entry will always exhibit a single trait combination, leading to HTI = 0 and therefore *µ*_*r*_ = 0, whereas a region with more entries has a positive probability of obtaining a non-zero HTI and therefore *µ*_*r*_ > 0. Since the classical CLT (Lindeberg–Lévy CLT) assumes that the random variables are iid, we turn to a different version of CLT that applies to independent, but not identically distributed, random variables – the Lyapunov CLT:

##### Lyapunov CLT

Let *X*_1_, *X*_2_, … *X*_*m*_ be independent random variables with 𝔼*X*_*i*_ = *µ*_*i*_ and 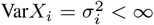. Denote *Y*_*i*_ = *X*_*i*_ − *µ*_*i*_.

(Lyapunov Condition) If there exists *δ >* 0 such that:

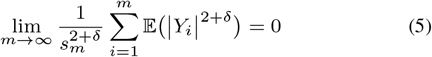

where:

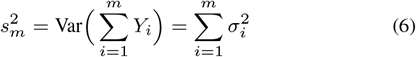

then:

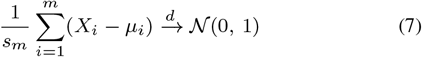

In our case, we want to use this theorem with 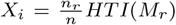 and *m* = *R* and obtain:

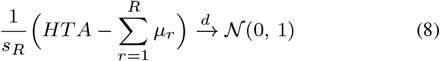

We observe that the Lyapunov condition is satisfied in our case. For any *δ >* 0,

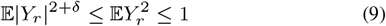

because *Y*_*r*_ ∈ [0, 1] (since HTI ∈ [0, 1]).

Therefore:

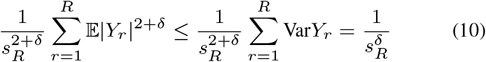

where the first inequality follows from Equation 9 combined with the fact that 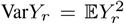 and the equality follows immediately from the definition of 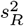 (Equation 6).

It remains to show that *s*_*R*_ → ∞ as *R* → ∞. Indeed, under the null hypothesis, the set of variances 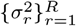 is bounded away from zero if we assume that there are no single-sample regions (otherwise we may increase the region size), or that there is a constant number of such regions.

Given a specific dataset, in order to use Lyapunov CLT, we must estimate *µ*_*r*_ and *σ*_*r*_ for all relevant (non-empty) regions, *r* = 1, …, *R*. We do so by simulating 1, 000 random-uniform permutations of the trait combination matrix (while holding constant the original positions of non-empty elements) and for each permutation we compute HTIs for all relevant regions. Then, for each region *r* ∈ *{*1, …, *R}*, we use its 1, 000 HTIs to estimate *µ*_*r*_ and *s*_*r*_.

We emphasize that the normal approximation holds only for sufficiently large values of *R*. We also note that for adequate interpretation of the HTA results, one should consider two one-sided *p*-values. Namely *p* and 1 − *p*, which represent the alternative hypotheses of homogeneity and heterogeneity, respectively. To determine the overall significance, the smaller *p*-value should be doubled.

## 3 Results

In this section we demonstrate the use of HTA in several domains. We begin with synthetic data for both 2-dimensional and 3-dimensional spatial data (Section 3.1). We then apply HTA to two 2-dimensional spatial transcriptomics datasets: Visium spatial RNA-seq by 10x Genomics (Section 3.2) and spatial transcriptomics inferred from pathology whole-slide images (Section 3.3). We also demonstrate a 3-dimensional use case using MRI images (Section 3.4). Finally, we demonstrate that HTA extends to other domains by analysing US census data (Section 3.5)

### 3.1 Synthetic data

In Figures 1 and 2, we depict results from applying HTA to 2-dimensional heterogeneity maps of shape (32, 32), each of which represents a trait-combination matrix. Regions are the visible square areas that fall between the grid lines. In Figure 1 we see heterogeneity maps for two traits, each of which has a random uniform distribution with probability 0.5 of occurrence in each position. As one would expect, the null hypothesis that the trait-combinations are randomly distributed is not rejected in all three region sizes (2, 8 and 16) since HTA *p*-values are > 0.3.

Figure 2 demonstrates that HTA discerns between homogeneous and heterogeneous distributions. Holding the region size constant, we observe that a perfectly homogeneous distribution within each region (left) is significantly homogeneous (HTA *p*-value *≅* 0), while a perfect heterogeneous distribution (right) is significantly heterogeneous (HTA 1-*p*-value *≅* 0). In comparison, a random heterogeneous distribution from *H*_0_ (middle) is neither (HTA *p*-value 0.5). This means that both significant homogeneity and significant heterogeneity are identified using HTA.

In Figure 5 we applied HTA to a 3-dimensional heterogeneity map of shape (32, 32, 3) (for the *x, y, z* axes, respectively). Since the region size can now also vary across the *z*-axis, we use two region sizes that differ only along this axis: (8, 8, 1) and (8, 8, 3) as illustrated at the bottom of Figure 5. Using a region size of (8, 8, 1), as in Figure 5 (left), where each region manifests exactly one of the trait combinations, we obtain significant homogeneity (HTA *p*-value *≅* 0). Conversely, using a region size of (8, 8, 3), illustrated in Figure 5 (right), where each region contains an equal amount of each of the three trait combinations, we obtain significant heterogeneity (HTA *p*-value of *≅* 1).

**Fig. 5.**
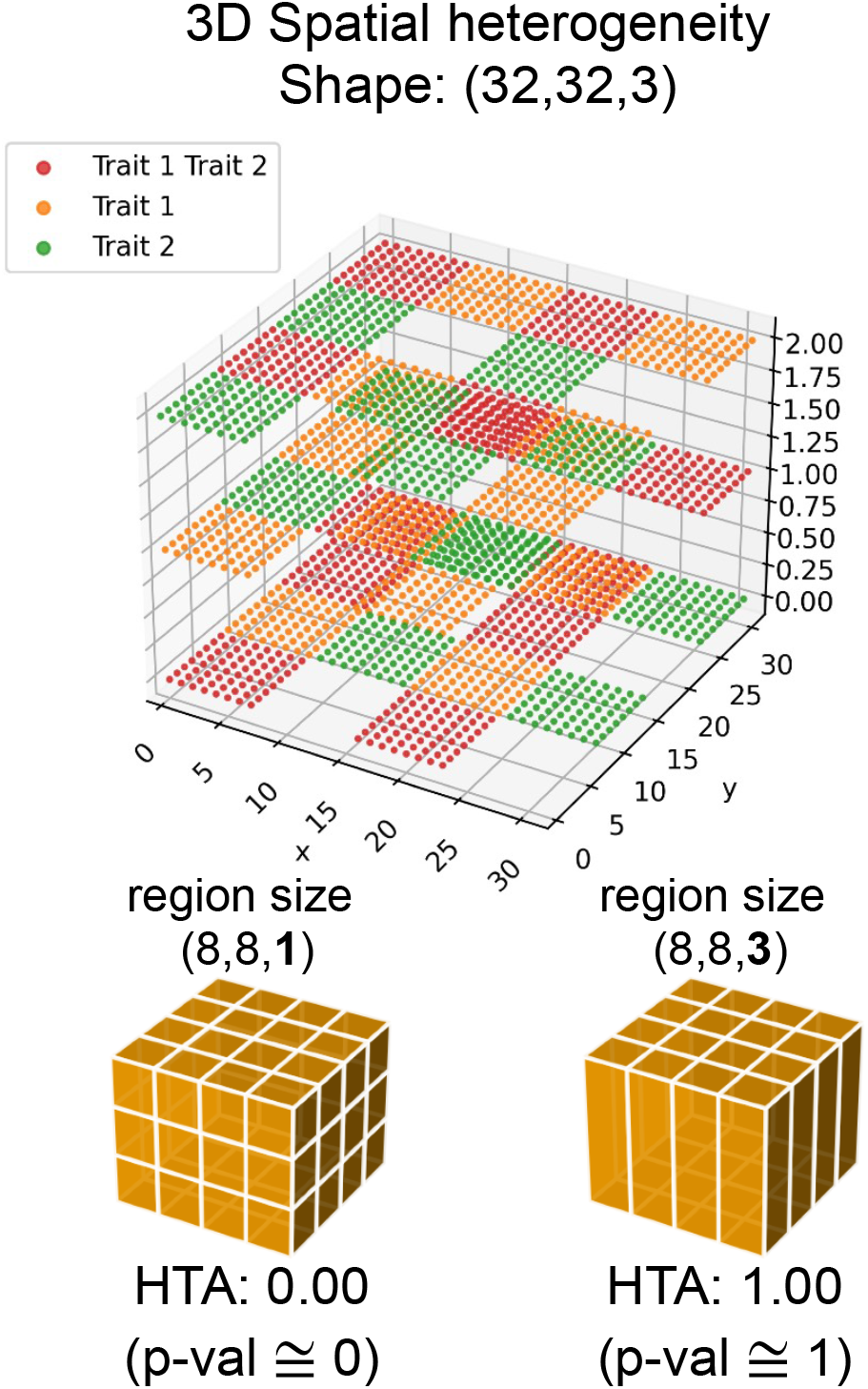
A 3D heterogeneity map and HTA for two region sizes that differ along the *z*-axis. The two illustrations below describe the resulting regions and are followed by HTA results: (left) region size (8, 8, 1) each region is perfectly homogeneous, yielding HTA 0 and HTA *p*-value *≅* 0; (right) region size (8, 8, 3) – each region is perfectly heterogeneous, yielding HTA 1 and HTA *p*-value *≅* 1.

### 3.2 Spatial RNA sequencing

We use 10x Genomics’ Visium breast cancer spatial gene-expression data (see Supplementary S2 for details). The sample is a Stage Group IIA breast cancer of type Luminal B (ER positive, PR negative and HER2 positive). To determine *n*_*rc*_, the number of entries in region *r* manifesting trait-combination *c*, we use the median threshold for each gene. An entry manifests combination *c* if all the genes in this combination are above their respective median expression level. Since this sample is Luminal B, it is expected that ESR1, FOXA1 and GATA3 would be spatially co-expressed [9], leading to a significantly homogeneous HTA index. We tested both two and three traits. Indeed, in Figure 3 B-D we observe that the tissue sample is significantly spatially homogeneous at both a local and global resolution (smaller and larger region-sizes, respectively), obtaining HTA *p*-values of < 10^−10^. This is compared to a random permutation of the observed trait combinations under *H*_0_ (Figure 3 A) which obtains an HTA *p*-value of 0.26 even at the smallest region size of 5. For the three traits: ESR1, GATA3 and FOXA1 (Figure 3 F-H) we observe similar results, with HTA *p*-value of < 10^−8^ at all three region sizes. This is in comparison to the random permutation of the observed trait combinations (Figure 3 E) which obtains an HTA *p*-value of 0.56 at region size 5.

Previous research has shown that T-cells remaining at the periphery of cancer cells, with low tumour infiltration, may be indicative of poor prognosis compared to tumours with high T-cell infiltration [16]. HTA can help identify such cases. In Figure 4 B-D we generated the heterogeneity map for ERBB2 (HER2) and CD8A (T-cells) using the same HER2 positive breast cancer sample. We focused on the area where ESR1 and GATA3 are relatively co-expressed (bottom half of the tissue in Figure 3) since, as explained above, these regions are likely to have high tumour content. Using HTA, we observe significant homogeneity in the two smaller region-sizes, 5 and 15, with HTA *p*-values < 10^−6^. Interestingly, at the larger region size of 30 we no longer observe significant homogeneity, with a *p*-value of 0.1. For context, a random dispersion of these T-cells (Figure 4A) is not significantly homogeneous, with an HTA *p*-value of 0.47. The significant homogeneity at the two smaller region sizes mean that the tumour cells are rarely, if at all, infiltrated by T-cells. However, the lack of significant homogeneity at the larger region size indicates that there is at least some infiltration of T-cells (otherwise we would observe significant homogeneity at this level too).

Since HTA is designed to handle a large number of traits, it is capable of capturing certain characteristics of clonal composition. We demonstrate this using the same breast cancer sample and 7 breast cancer driver genes: MYC, ESR1, ERBB2, GATA3, FOXA1, TP53 and CDK4. In Figure 6 A we can see the resulting heterogeneity map. Using a region size of 15, we obtain a significantly homogeneous HTA (*p*-value: 10^−16^). In B we observe a random permutation of the observed trait combinations, which results in a non-significantly homogeneous HTA (*p*-value: 0.43). Since 7 traits give rise to 2^7^ − 1 = 127 non-empty combinations (provided that all are present), we do not attempt to display the legend. Instead, we produce a ‘region-report’ to identify the most frequent combinations in regions of interest. For example, in the bottom left corner, at (−0.5, 59.5), the most frequent combination is all 7 driver genes, accounting for 73% of the elements in that region. The second most frequent combination is the 6 driver genes that remain after removing CDK4. While the region to its right has a similar composition, two regions to the right, at (29.5, 59.5), already exhibits a different, yet relatively homogeneous composition: the combination of all 7 account for 41% of the elements; a small number of different combinations of 6 of the driver genes account for 26% and; the vast majority of the remaining elements (accounting for 28%) are several combinations of 4 and 5 genes. These observations align with the significantly homogeneous HTA, and may indicate that the significant homogeneity may be due to the gradual formation of a dominant subclone that over-expresses all 7 genes.

**Fig. 6.**
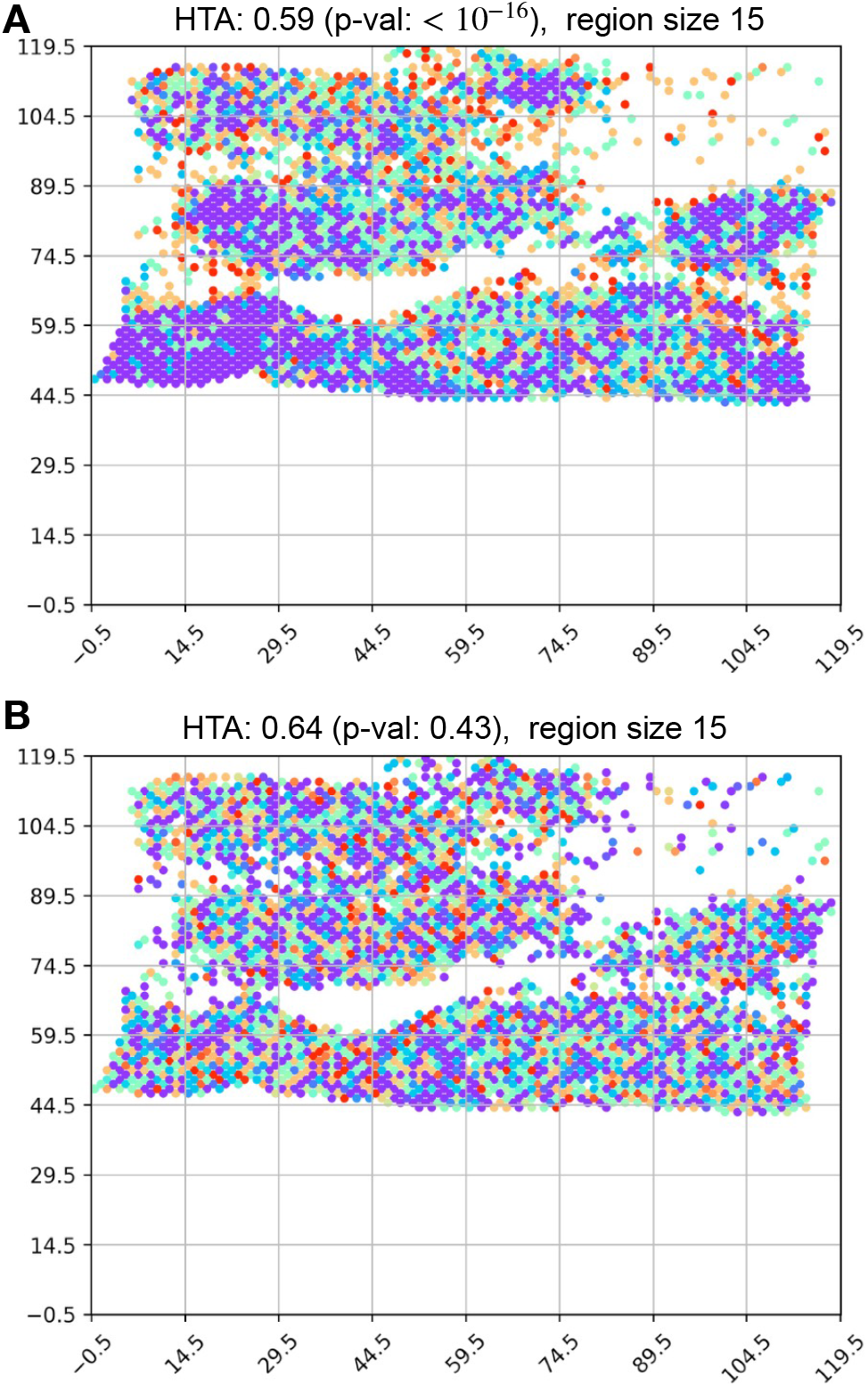
Heterogeneity maps and HTA for 7 breast cancer driver genes: MYC, ESR1, ERBB2, GATA3, FOXA1, TP53 and CDK4. (A) actual heterogeneity map, with HTA *p*-value of 10^−16^ at region size 15; (B) heterogeneity map under *H*_0_ (random permutation of the trait-combinations), with HTA *p*-value of 0.43.

### 3.3 Spatial transcriptomics from pathology whole-slide images

In this section we use spatial transcriptomics inferred from breast cancer pathology whole-slide images obtained from [10]. This dataset contains 324 subjects for which: (a) MKI67 and miR-17 expression were spatially resolved to their respective pathology whole-slide images, yielding binary maps that indicate where each gene was detected as over-expressed; and (b) survival data is available. Since not all slides, and therefore inferred maps, have the same size, we first resize them to (90, 90), chosen based on their median shape. To ensure that the maps remain binary, we use nearest neighbour interpolation during resizing. We apply HTA using the three region sizes: 15, 30 and 45. We then split the cohort into two equal sets based on their HTAs: > median HTA and ≤ median HTA (relatively heterogeneous and relatively homogeneous, respectively). Figure 7 shows the results for the survival analysis performed using these heterogeneity-based assignments. We observe significant survival differences in the two larger region sizes, with the middle one, region size 30, obtaining the lowest *p*-value of 0.01.

**Fig. 7.**
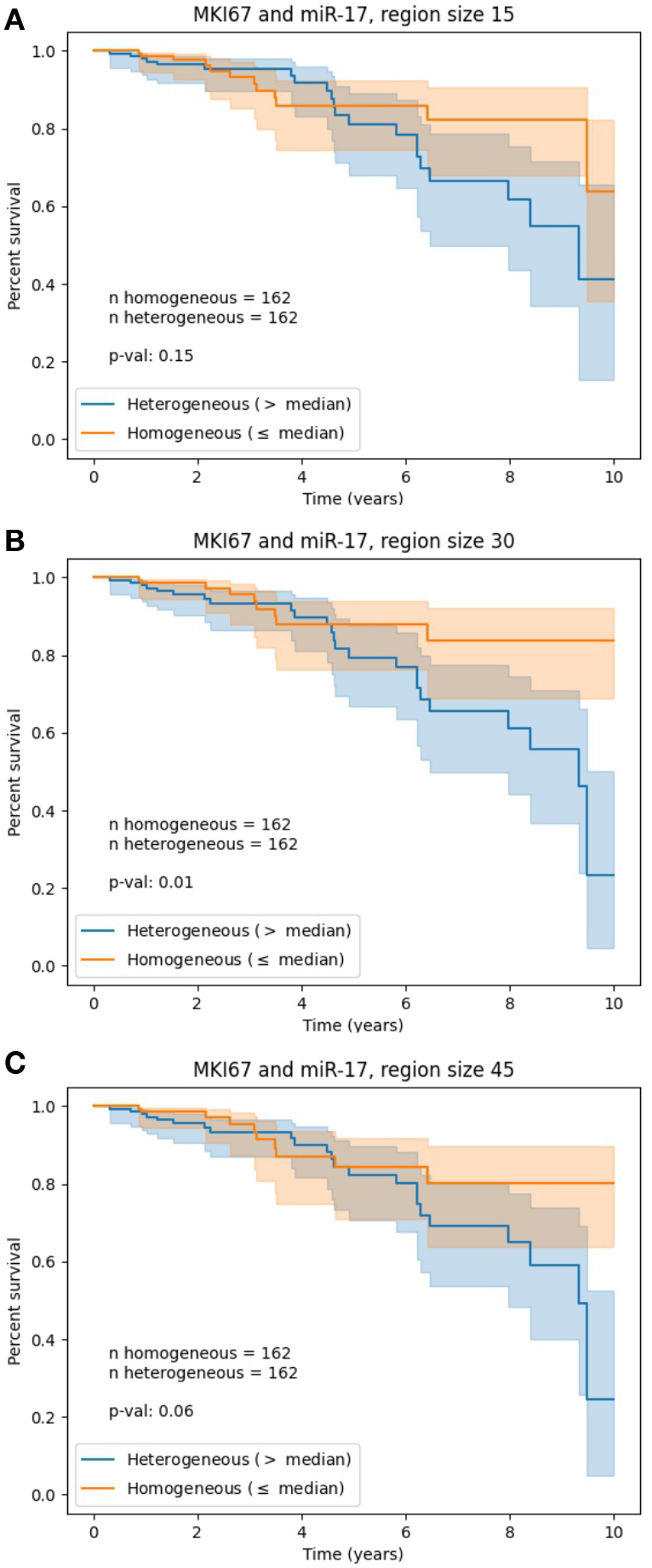
Survival analysis with respect to HTA derived from two spatially-resolved traits in breast cancer pathology whole-slides – MKI67 and miR-17 expression level. Binary spatial transcriptomics maps, inferred from the slides of 324 subjects, were split into high and low HTA with respect to the cohort’s median: > median (blue) and ≤ median (orange). The plots differ in the region size used to compute HTA. HTA region sizes and corresponding log-rank *p*-values: (A) 15, *p* = 0.15; (B) 30, *p* = 0.01; (C) 45, *p* = 0.06. All maps were resized to (90, 90), close to the median map size, using nearest neighbour interpolation. Each survival curve is shown with a 95% confidence interval.

### 3.4 MRI

In this section we demonstrate the use of HTA in the context of 3D data by analysing brain MRI scans (see Supplementary S3 for further details). We use 4 axial MRI scans from 4 different subjects, representing: (1) normal ageing, (2) Alzheimer’s disease (3) Metastatic bronchogenic carcinoma and (4) Glioma. For each subject, we obtained the only two weighted sequences that were available for all four: T2-weighted and Proton density (PD) weighted (these accentuate different properties; for details see, e.g., [4]). We use these as the traits, where a strong signal (brighter) gets 1 and a low signal (darker) gets 0, depending on whether they are above or below the median grayscale value, correspondingly. Since MRI scans are sequences of images (slices), they represent a 3-dimensional space. We use 3 different slices per subject taken from similar locations in each (Supplementary S3). We use 3 for visualisation purposes, but the analysis applies to any number of slices. In Figure 8 we see the resulting 3D heterogeneity maps and HTA for all four subjects. The shape of each map is (256, 256, 3) and the region size is (128, 128, 3). Normal ageing (top) is the only one that is not significantly homogeneous at a 0.01 threshold (*p*-value 0.02). The remaining three, each of which represents a different disease, are significantly homogeneous. The figures are ordered in decreasing *p*-values. Interestingly, the two cancer scans obtain the strongest significance, with *p*-values 10^−84^ (metastatic bronchogenic carcinoma) and 10^−149^ (glioma). Alzheimer’s disease is also significantly homogeneous, with a *p*-value of 0.0002.

**Fig. 8.**
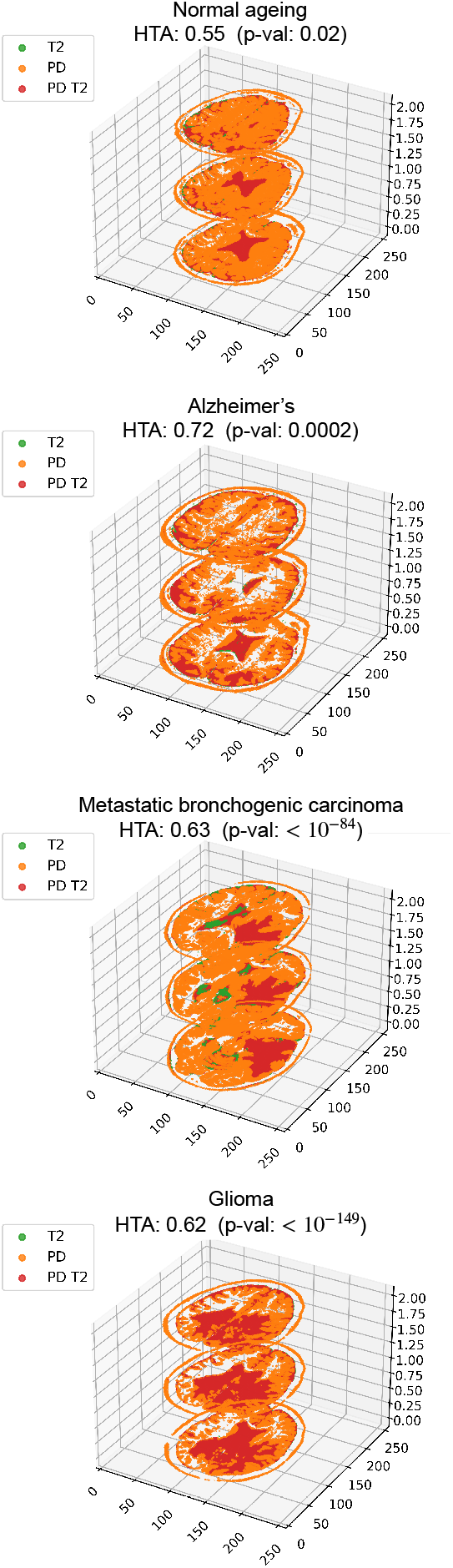
Heterogeneity maps and HTA for 4 MRI scans taken from different subjects. The shape of each heterogeneity map is (256, 256, 3) and the region size is (128, 128, 3). From top to bottom (highest to lowest HTA *p*-values): (1) normal ageing, (2) Alzheimer’s disease (3) Metastatic bronchogenic carcinoma and (4) Glioma.

We note that for images, there may be other binarization techniques besides the median that are worth considering (e.g., Li thresholding to detect background vs foreground). Specifically for MRI, it may also be relevant to use actual raw signal measurements, provided such data is available. In such a case, other binarization methods may be better suited than the median value threshold. We have kept the median threshold for simplicity of demonstration.

### 3.5 Census data

To demonstrate the overall utility of HTA we also applied it to recent census data for 3, 092 counties across the US, excluding those in Alaska, Hawaii and Puerto Rico due to their relatively large distance from the other states (see Supplementary S5). For each county, the data contains the total population per ethnicity, across multiple ethnicity groups. In Figure 9 we observe the heterogeneity map and corresponding HTA index and *p*-value obtained for the three ethnicity-related traits: “Black or African American alone > 5% of total population in county”, “Asian alone > 5% of total population in county” and “American Indian and Alaska Native alone > 5% of total population in county” for region size 70. We observe significant homogeneity, with HTA *p*-value 10^−19^, indicating that people from the same ethnic origin tend to cluster in specific regions. Since all regions included over 5% “White alone” we did not include this category.

**Fig. 9.**
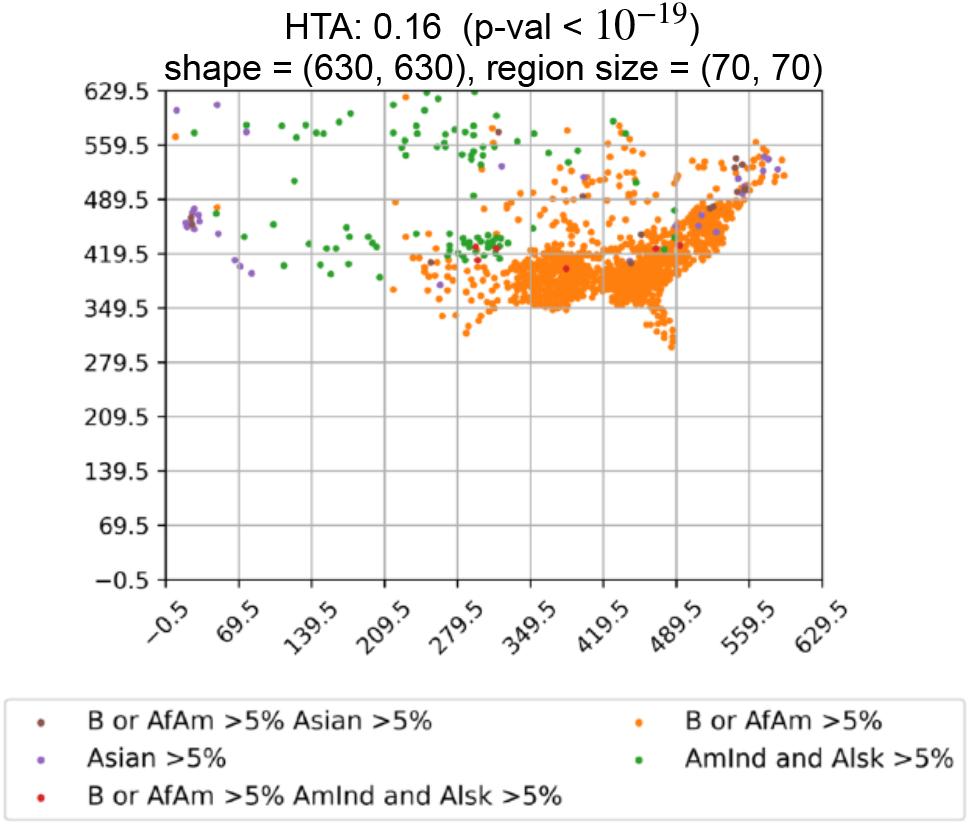
Heterogeneity maps and HTA for US census data for three ethnicity-related traits: “Black or African American alone > 5% of total population in county”, “Asian alone > 5% of total population in county” and “American Indian and Alaska Native alone > 5% of total population in county” using region size 70. We observe significant homogeneity with HTA *p*-value < 10^−19^.

Since the size of such maps could potentially be much larger, we also tested whether 100 random-uniform permutations (required to compute Lyapunov CLT parameters, as described in Section 2.3.2), instead of 1, 000, would be sufficient to obtain similar HTAs and *p*-values. The results for 100 repeats are nearly identical to those of 1, 000 repeats: we obtain *p*-values 10^−19^ for 1, 000 and 10^−18^ for 100.

## 4 Discussion

HTA provides a solution to the growing need for statistical analysis tools that are capable of quantifying spatial heterogeneity in the complex setting of high throughput molecular biology data, including spatial transcriptomics and digital pathology. It is also useful in other domains, including imaging and geographic information systems. HTA accurately reflects spatial heterogeneity at multiple resolutions, can handle a large number of variables (trait-combinations) and lends itself to efficient statistical assessment. In the context of addressing multiple traits, we also note the difference between HTA and the existing literature. For example, Figure 10 demonstrates that Morisita-Horn, which measures the overlap between two traits across all regions of a given space, will declare a perfect overlap for both examples shown in the figure, whereas HTA successfully discerns between the two.

While HTA is multivariate, it may not easily scale to hundreds or thousands of molecular traits. This can be overcome by using aggregation methods to obtain several meta-traits, each representing a group of individual traits. We demonstrate this in Supplementary S6 using immune pathway enrichment scores (computed using GSVA [**?**]), representing a total of 69 genes, and using cluster IDs obtained from 10X’s Loupe Browser, representing 2, 786 genes. Using such aggregation methods HTA can be applied to data that spans a large number of individual traits.

We have also shown that HTA can be used at multiple scales, controlled by the region-size parameter. We note that other multi-scale measures, such as 2D multi-scale entropy (e.g., [**?**]), as well as fractal-based and wavelet-based methods (e.g., [**?**]), while multi-scale, do not apply to the trait-combination representation that HTA can analyse. These methods apply either to 2D images, with a strong emphasis on relationships between colors, or to 2D binary matrices. Since the colors in the heterogeneity maps are not ordinal, and since there is more than one trait-combination, neither option is relevant.

HTA also has other advantages. First, HTA is simple to use since it requires a single input parameter (region size) and easy to interpret since it is directly derived from Shannon’s entropy. This is in contrast to available tools that are limited in at least one of these aspects. HTA also applies to any d-dimensional space. We demonstrated this in 3-dimensions using both synthetic and MRI data. Finally, HTA extends beyond the scope of molecular biology and medical imaging and can be used in many other domains, as was demonstrated with US census data.

HTA can be further used in down-stream analyses. For example, in the case of ERBB2 (HER2) and CD8A (T-cells) demonstrated in Figure 4, different HTAs could potentially be associated with different responses to immune therapy treatment. Results of applying HTA in digital pathology show that HTA may be predictive of survival in breast cancer.

HTA offers many potential extensions worthy of investigation in future work. One option is to combine HTA scores from multiple scales into a new measure that summarizes them. It may also be relevant to extend HTA to apply to continuous measurements. Finally, alternative permutation-based null models may also be investigated. For example, a locality preserving null model (where the region can be determined by some radius) may be useful in certain cases where retaining local characteristics is important. We note, however, that it may not be appropriate when considering more global phenomenons. One such case is when measuring the level of T-cell infiltration, as demonstrated in Figure 4; had we performed a locality preserving permutation, we would not have been able to assess the significance of infiltration since such a permutation would cause the null model to assume that there *is* T-cell infiltration to begin with (this sample has immune cells around cancer cells in most locations). Nevertheless, alternative null models may be highly relevant in other cases, and offer interesting opportunities for further investigation.

As spatial transcriptomics data and digital pathology inference techniques become increasingly available and accurate, we expect methods that address spatial distributions, including HTA and its potential extensions, to become ubiquitous.

## Supporting information

HTA Supplementary

## Acknowledgements

We thank the Technion Computer Science Department for generously supporting ALJ. This project received funding from the European Union’s Horizon 2020 Research and Innovation Programme under Grant Agreement No. 847912. We thank the Yakhini Research Group and Vessela N. Kristensen for important discussions and input.

## Author contributions

ALJ and ZY conceived the idea and developed the approach. ALJ developed and wrote the software package. XT advised on breast cancer molecular biology.

## References

[1] 10x Genomics - Spatial Transcriptomics. https://www.10xgenomics.com/spatial-transcriptomics/. Accessed: 2020-12-24.

[2] Khalid AbdulJabbar, Shan E Ahmed Raza, Rachel Rosenthal, Mariam Jamal-Hanjani, Selvaraju Veeriah, Ayse Akarca, Tom Lund, David A Moore, Roberto Salgado, Maise Al Bakir, et al. Geospatial immune variability illuminates differential evolution of lung adenocarcinoma. Nature Medicine, pp. 1–9, 2020.

[3] Alma EV Andersson, Ludvig Larsson, Linnea Stenbeck, Fredrik Salmén, Anna Ehinger, Sunny Z Wu, Ghamdan Al-Eryani, Daniel L Roden, Alexander Swarbrick, Ake Borg, et al. Spatial deconvolution of her2-positive breast tumors reveals novel intercellular relationships. bioRxiv, 2020.

[4] AL Chong, Ronil V Chandra, KC Chuah, E. Roberts, and SL Stuckey. Proton density mri increases detection of cervical spinal cord multiple sclerosis lesions compared with t2-weighted fast spin-echo. American Journal of Neuroradiology, 37(1):180–184, 2016.

[5] Nicolas Coudray, Paolo Santiago Ocampo, Theodore Sakellaropoulos, Navneet Narula, Matija Snuderl, David Fenyö, Andre L Moreira, Narges Razavian, and Aristotelis Tsirigos. Classification and mutation prediction from non–small cell lung cancer histopathology images using deep learning. Nature medicine, 24(10):1559–1567, 2018.

[6] Stephanie M Dobson, Laura García-Prat, Robert J Vanner, Jeffrey Wintersinger, Esmé Waanders, Zhaohui Gu, Jessica McLeod, Olga I Gan, Ildiko Grandal, Debbie Payne-Turner, et al. Relapse-fated latent diagnosis subclones in acute b lineage leukemia are drug tolerant and possess distinct metabolic programs. Cancer Discovery, 10(4):568–587, 2020.

[7] Robert J Gillies, Daniel Verduzco, and Robert A Gatenby. Evolutionary dynamics of carcinogenesis and why targeted therapy does not work. Nature Reviews Cancer, 12(7):487–493, 2012.

[8] Shona Hendry, Roberto Salgado, Thomas Gevaert, Prudence A Russell, Tom John, Bibhusal Thapa, Michael Christie, Koen Van De Vijver, M Valeria Estrada, Paula I Gonzalez-Ericsson, et al. Assessing tumor infiltrating lymphocytes in solid tumors: a practical review for pathologists and proposal for a standardized method from the international immuno-oncology biomarkers working group: Part 1: Assessing the host immune response, tils in invasive breast carcinoma and ductal carcinoma in situ, metastatic tumor deposits and areas for further research. Advances in anatomic pathology, 24(5):235, 2017.

[9] Jocelyne Jacquemier, Emmanuelle Charafe-Jauffret, Florence Monville, Benjamin Esterni, Jean Marc Extra, Gilles Houvenaeghel, Luc Xerri, François Bertucci, and Daniel Birnbaum. Association of gata3, p53, ki67 status and vascular peritumoral invasion are strongly prognostic in luminal breast cancer. Breast Cancer Research, 11(2):R23, 2009.

[10] Alona Levy-Jurgenson, Xavier Tekpli, Vessela N Kristensen, and Zohar Yakhini. Spatial transcriptomics inferred from pathology whole-slide images links tumor heterogeneity to survival in breast and lung cancer. Scientific reports, 10(1):1–11, 2020.

[11] Suzanne E Little, Sergey Popov, Alexa Jury, Dorine A Bax, Lawrence Doey, Safa Al-Sarraj, Juliane M Jurgensmeier, and Chris Jones. Receptor tyrosine kinase genes amplified in glioblastoma exhibit a mutual exclusivity in variable proportions reflective of individual tumor heterogeneity. Cancer research, 72(7):1614–1620, 2012.

[12] Fei Ma, Yanfang Guan, Zongbi Yi, Lianpeng Chang, Qiao Li, Shanshan Chen, Wenjie Zhu, Xiuwen Guan, Chunxiao Li, Haili Qian, et al. Assessing tumor heterogeneity using ctdna to predict and monitor therapeutic response in metastatic breast cancer. International Journal of Cancer, 146(5):1359–1368, 2020.

[13] Carlo C Maley, Konrad Koelble, Rachael Natrajan, Athena Aktipis, and Yinyin Yuan. An ecological measure of immune-cancer colocalization as a prognostic factor for breast cancer. Breast Cancer Research, 17(1):131, 2015.

[14] Takahiro Masuda, Roman Sankowski, Ori Staszewski, Chotima Böttcher, Lukas Amann, Christian Scheiwe, Stefan Nessler, Patrik Kunz, Geert van Loo, Volker Arnd Coenen, et al. Spatial and temporal heterogeneity of mouse and human microglia at single-cell resolution. Nature, 566(7744):388–392, 2019.

[15] Ines P Nearchou, Bethany M Gwyther, Elena CT Georgiakakis, Christos G Gavriel, Kate Lillard, Yoshiki Kajiwara, Hideki Ueno, David J Harrison, and Peter D Caie. Spatial immune profiling of the colorectal tumor microenvironment predicts good outcome in stage ii patients. NPJ Digital Medicine, 3(1):1–10, 2020.

[16] Giancarlo Pruneri, Andrea Vingiani, and Carsten Denkert. Tumor infiltrating lymphocytes in early breast cancer. The Breast, 37:207–214, 2018.

[17] Grzegorz A Rempala and Michal Seweryn. Methods for diversity and overlap analysis in t-cell receptor populations. Journal of mathematical biology, 67(6-7):1339–1368, 2013.

[18] Brian D Ripley. The second-order analysis of stationary point processes. Journal of applied probability, 13(2):255–266, 1976.

[19] Jin-Feng Wang, Tong-Lin Zhang, and Bo-Jie Fu. A measure of spatial stratified heterogeneity. Ecological Indicators, 67:250–256, 2016.

[20] Yinyin Yuan. Spatial heterogeneity in the tumor microenvironment. Cold Spring Harbor perspectives in medicine, 6(8):a026583, 2016.

[21] Xiang Zheng, Andreas Weigert, Simone Reu, Stefan Guenther, Siavash Mansouri, Birgit Bassaly, Stefan Gattenlöhner, Friedrich Grimminger, Soni Savai Pullamsetti, Werner Seeger, et al. Spatial density and distribution of tumor-associated macrophages predict survival in non–small cell lung carcinoma. Cancer Research, 80(20):4414–4425, 2020.

