## Supplementary material for "Assessing heterogeneity in spatial data using the HTA index with applications to spatial transcriptomics and imaging": HTA Supplementary

*Bioinformatics*
doi.10.1093/bioinformatics/xxxxxx
Advance Access Publication Date: Day Month Year
Manuscript Category

OXFORD

Subject Section

### Supplementary Material: Assessing heterogeneity in spatial data using the HTA index with applications to spatial transcriptomics and imaging

**Alona Levy-Jurgenson [1,*], Xavier Tekpli [2,3] and Zohar Yakhini [1,4*]**

[1] Department of Computer Science, Technion - Israel Institute of Technology, Haifa 32000, Israel
[2] Department of Medical Genetics, Institute of Clinical Medicine, University of Oslo and Oslo University Hospital, Oslo, Norway
[3] Department of Cancer Genetics, Institute for Cancer Research, Oslo University Hospital, 0310 Oslo, Norway and
[4] Arazi School of Computer Science, Interdisciplinary Center, Herzliya 4610101, Israel.

*To whom correspondence should be addressed.

Associate Editor: XXXXXXX

Received on XXXXX; revised on XXXXX; accepted on XXXXX

#### Abstract

**Motivation:** Tumour heterogeneity is being increasingly recognised as an important characteristic of cancer and as a determinant of prognosis and treatment outcome. Emerging spatial transcriptomics data hold the potential to further our understanding of tumour heterogeneity and its implications. However, existing statistical tools are not sufficiently powerful to capture heterogeneity in the complex setting of spatial molecular biology.

**Results:** We provide a statistical solution, the HeTerogeneity Average index (HTA), specifically designed to handle the multivariate nature of spatial transcriptomics. We prove that HTA has an approximately normal distribution, therefore lending itself to efficient statistical assessment and inference. We first demonstrate that HTA accurately reflects the level of heterogeneity in simulated data. We then use HTA to analyse heterogeneity in two cancer spatial transcriptomics datasets: spatial RNA sequencing by 10x Genomics and spatial transcriptomics inferred from H&E. Finally, we demonstrate that HTA also applies to 3D spatial data using brain MRI. In spatial RNA sequencing we use a known combination of molecular traits to assert that HTA aligns with the expected outcome for this combination. We also show that HTA captures immune-cell infiltration at multiple resolutions. In digital pathology we show how HTA can be used in survival analysis and demonstrate that high levels of heterogeneity may be linked to poor survival. In brain MRI we show that HTA differentiates between normal ageing, Alzheimer's disease and two tumours. HTA also extends beyond molecular biology and medical imaging, and can be applied to many domains, including GIS.

#### 1 Supplementary Material

##### 1.1 Supplementary Note S1 - HTA Monotonicity Proof

In this section we prove that HTA monotonically decreases with grid refinement. We first provide several definitions: regions of a trait-combination matrix, $HTI$ and $HTA$. We begin the proof by showing that $HTA \leq HTI$ for any given trait-combination matrix. We then use this result along with Gibbs' inequality to prove that $HTA$ monotonically decreases with grid refinement.

###### 1.1.1 Definitions

Let $M$ be a matrix. Let $C$ be the number of non-empty trait combinations that may be observed in $M$. Let the value of cell $(i, j)$ in $M$ indicate which trait combination is present at the corresponding spatial location in the sample (or 'none' otherwise). Then we call $M$ a trait-combination matrix. For example, $M$'s entries may each represent a single barcode in spatial RNA-sequencing or a single tile in digital pathology.

© The Author 2015. Published by Oxford University Press. All rights reserved. For permissions, please

*Definition (region of matrix)* Denote by $M\big|_G$ the set of non-empty (not all 'none') sub-matrices obtained by applying grid $G$ on $M$:
$M\big|_G = \{M_1, M_2, ..., M_R\}$, where $R$ is the total number of non-empty sub-matrices. Then the *region* $M_r$ of $M$ is the $r$-th non-empty sub-matrix in $M\big|_G$. The number of entries in region $M_r$ that are not 'none' is denoted by $n_r$.

*Definition (HTI)* We will mainly use the following definition of HTI, which, as explained below, is equivalent to the definition provided in the main text:

$$HTI(M) := -\sum_{r=1}^{R}\sum_{c=1}^{C}\frac{n_{rc}}{n}\log\sum_{r=1}^{R}\frac{n_{rc}}{n} \qquad (1)$$

where $n_{rc}$ is the number of cells $(i, j) \in M_r$ indicating combination $c$, and $n$ is the total number of cells $(i, j) \in M$ with at least one trait. Note that $\sum_{c=1}^{C}\frac{n_{rc}}{n_r} = 1$.

To see that this is equivalent to the definition of HTI provided in the main text:

$$HTI = -\sum_{c=1}^{C} q_c \log_C(q_c) \qquad (2)$$

we note that $q_c$, the proportion of spatial positions for which exactly all traits in combination $c$ manifest, can be re-written as a function of the number of samples observed in each region:

$$q_c = \sum_{r=1}^{R}\frac{n_{rc}}{n}$$

which give us:

$$HTI(M) = -\sum_{c=1}^{C}\sum_{r=1}^{R}\frac{n_{rc}}{n}\log\sum_{r=1}^{R}\frac{n_{rc}}{n}$$

and we can interchange the sums to obtain Equation 1.

*Definition (HTA)* Let $M\big|_G = \{M_1, M_2, ..., M_R\}$ be the set of regions obtained by applying grid $G$ on some trait-combination matrix $M$. Let $\{n_1, n_2, ..., n_R\}$ be the corresponding number of entries in each region that are not 'none'. Then we define:

$$HTA(M\big|_G) := \sum_{r=1}^{R}\frac{n_r}{n}HTI(M_r) = \sum_{r=1}^{R}\frac{n_r}{n}\left(-\sum_{c=1}^{C}\frac{n_{rc}}{n_r}\log\frac{n_{rc}}{n_r}\right) \qquad (3)$$

where the last equality follows the definition of HTI in Equation 2, when applied to the matrix $M_r$.

##### 1.1.2 Proving HTA monotonically decreases with grid refinement

Note that this is similar to the fact that:

$$H(Y|X) \leq H(Y)$$

for random variables $X$ and $Y$.

Below we provide a proof that specifically applies to our case. We first prove that $HTA(M\big|_G) \leq HTI(M)$ and then use this to prove that HTA monotonically decreases with grid refinement.

**Proving $HTA(M\big|_G) \leq HTI(M)$**

The proof uses Gibb's inequality, which states the following:

Let $P = p_1, p_2, \ldots, p_n$ and $Q = q_1, q_2, \ldots, q_n$ be two different probability distributions. Then:

$$-\sum_{i=1}^{n} p_i \log p_i \leq -\sum_{i=1}^{n} p_i \log q_i$$

**Proposition 1.** $HTA(M\big|_G) \leq HTI(M)$

We would like to show that:

$$\sum_{r=1}^{R}\frac{n_r}{n}\left(-\sum_{c=1}^{C}\frac{n_{rc}}{n_r}\log\frac{n_{rc}}{n_r}\right) \leq -\sum_{r=1}^{R}\sum_{c=1}^{C}\frac{n_{rc}}{n}\log\sum_{r=1}^{R}\frac{n_{rc}}{n}$$

Proof. If the inequality holds for each of the $r$ terms individually, then we are done. Therefore, it is sufficient to show that for all $r$ we have:

$$\frac{n_r}{n}\left(-\sum_{c=1}^{C}\frac{n_{rc}}{n_r}\log\frac{n_{rc}}{n_r}\right) \leq -\sum_{c=1}^{C}\frac{n_{rc}}{n}\log\sum_{r=1}^{R}\frac{n_{rc}}{n}$$

Multiplying by $\frac{n}{n_r}$ we can see that this is true iff:

$$-\sum_{c=1}^{C}\frac{n_{rc}}{n_r}\log\frac{n_{rc}}{n_r} \leq \left(-\sum_{c=1}^{C}\frac{n_{rc}}{n}\log\sum_{r=1}^{R}\frac{n_{rc}}{n}\right)\frac{n}{n_r}$$

(note that since empty regions are removed $n_r \neq 0$).
which give us:

$$-\sum_{c=1}^{C}\frac{n_{rc}}{n_r}\log\frac{n_{rc}}{n_r} \leq \sum_{c=1}^{C}\frac{n_{rc}}{n_r}\log\sum_{r=1}^{R}\frac{n_{rc}}{n}$$

Setting:

$$p_c = \frac{n_{rc}}{n_r} \qquad q_c = \sum_{r=1}^{R}\frac{n_{rc}}{n}$$

we can see Gibbs' conditions hold (see Lemma 1.2 to see why $q_c$ constitute a probability distribution), and therefore so does Gibbs' inequality. Therefore:
$HTA(M\big|_G) \leq HTI(M)$.

##### 1.1.3 Proving HTA monotonically decreases with grid refinement

**Definition 1.1.** (Finer grid)

Let $G$ and $G^f$ be two grids on $M$. We say that $G^f$ is *finer* than $G$ if for all $r$, $M_r \in G$ is precisely the union of regions $\{M_{rk}\}_{k=1...K} \in G^f$. In this case we write: $G^f < G$.

Theorem 1.1. *(HTA monotonically decreases with grid refinement)*
*Let G and $G^f$ be two grids on M, where $G^f \leq G$. Then:*

$$HTA(M\big|_{G^f}) \leq HTA(M\big|_G)$$

Proof. Let $M_r$ be a region in $G$ and let $\{M_{rk}\}_{k=1...K}$ be the set of regions in $G^f$ whose union is precisely $M_r$. Since $M_r$ is a also a matrix, we can view $\{M_{rk}\}_{k=1...K}$ as regions of $M_r$ obtained by applying the relevant part of the grid $G^f$ to $M_r$, denoted $G^f_r$.

Then we know from Proposition 1 that:

$$HTA(M_r\big|_{G^f_r}) \leq HTI(M_r)$$

But this is true for all regions $M_r$. Therefore this still holds if we multiply by $\frac{n_r}{n}$ (a constant per $r$) and sum over all regions:

$$\sum_{r=1}^{R}\frac{n_r}{n}HTA(M_r\big|_{G^f_r}) \leq \sum_{r=1}^{R}\frac{n_r}{n}HTI(M_r) \qquad (4)$$

Expanding the left term we obtain:

$$
\begin{aligned}
\sum_{r=1}^{R}\frac{n_r}{n}HTA(M_r|_{G_r^f}) &= \sum_{r=1}^{R}\frac{n_r}{n}\sum_{k=1}^{K}\frac{n_{rk}}{n_r}HTI(M_{rk})\\
&= \sum_{r=1}^{R}\sum_{k=1}^{K}\frac{n_{rk}}{n}HTI(M_{rk})\\
&= HTA(M|_{Gf})
\end{aligned}
$$

where the last equality is since we are summing over all regions in $M|_{Gf}$.

The right term of Eq. 4 is simply $HTA(M|_G)$, by definition. Combining this observation with the above expansion of the left term, Eq. 4 becomes:

$$HTA(M|_{Gf}) \leq HTA(M|_G)$$

Lemma 1.2. *(Gibbs' conditions hold)*

Proof. We set:

$$p_c = \frac{n_{rc}}{n_r} \qquad q_c = \sum_{r=1}^{R}\frac{n_{rc}}{n}$$

We know that $\sum_{c=1}^{C} p_c = 1$ (since $\sum_{c=1}^{C} n_{rc} = n_r$). It remains to show: $\sum_{c=1}^{C} q_c = 1$.

$$\sum_{c=1}^{C} q_c = \sum_{c=1}^{C}\sum_{r=1}^{R}\frac{n_{rc}}{n} = \sum_{r=1}^{R}\sum_{c=1}^{C}\frac{n_{rc}}{n} = \sum_{r=1}^{R}\frac{n_r}{n} = 1$$

#### 1.2 Supplementary Note S2 - 10x Visium Data

- Sample identifier: Human Breast Cancer (Block A Section 1) Spatial Gene Expression Dataset by Space Ranger 1.1.0
- Link: https://support.10xgenomics.com/spatial-gene-expression/datasets/1.1.0/V1_Breast_Cancer_Block_A_Section_1
- Files:
  "Feature / cell matrix (filtered)"
  "Spatial imaging data"

#### 1.3 Supplementary Note S3 - MRI

All MRI slices, such as those observed in Figure 1 were obtained from The Human Brain Atlas from the following links and slice numbers, for both PD-weighted and T2-weighted sequences. Note that 'Metastatic bronchogenic carcinoma' has different slice numbers since slice numbers do not necessarily correspond to the same brain regions in different subjects.

- Normal ageing:
  http://www.med.harvard.edu/aanlib/cases/case36/mr1/029.html
  Slices: 32,34,36
- Alzheimer's disease:
  http://www.med.harvard.edu/aanlib/cases/case40/mr2/041.html
  Slices: 32,34,36
- Glioma:
  http://www.med.harvard.edu/aanlib/cases/case1/mr1/026.html
  Slices: 32,34,36
- Metastatic bronchogenic carcinoma:
  http://www.med.harvard.edu/aanlib/cases/case28/mr2/013.html
  Slices: 09,11,13.

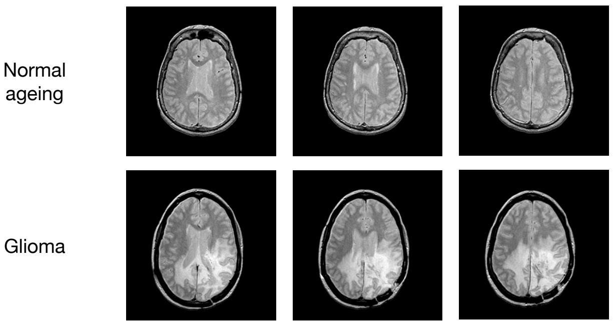


**Fig. 1.** An example of the raw MRI slices. This figure shows the three PD-weighted slices used in the analysis of normal ageing and glioma.

#### 1.4 Supplementary Note S4

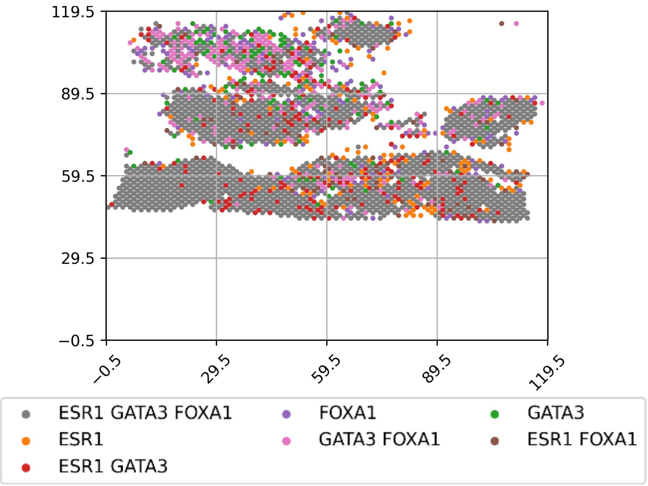


**Fig. 2.** Heterogeneity map for the three traits: ESR1, GATA3 and FOXA1, along with the full legend. Each color represents the manifestation of a different trait combination.

#### 1.5 Supplementary Note S5

US census data was obtained from the US Census Bureau website, under: Race and Ethnicity - County:

`https://covid19.census.gov/datasets/ace8fa8bea514d07a3139e4657b3cd9c_0`

No pre-processing was applied.

#### 1.6 Supplementary Note S6

**List of genes per pathway**
List of genes per pathway can be obtained from:

- https://www.gsea-msigdb.org/gsea/msigdb/cards/<NAME> where NAME is one of the aforementioned pathway names.

**GSVA package**
GSVA package documentation can be found at:

- https://www.bioconductor.org/packages/release/bioc/vignettes/GSVA/inst/doc/GSVA.pdf

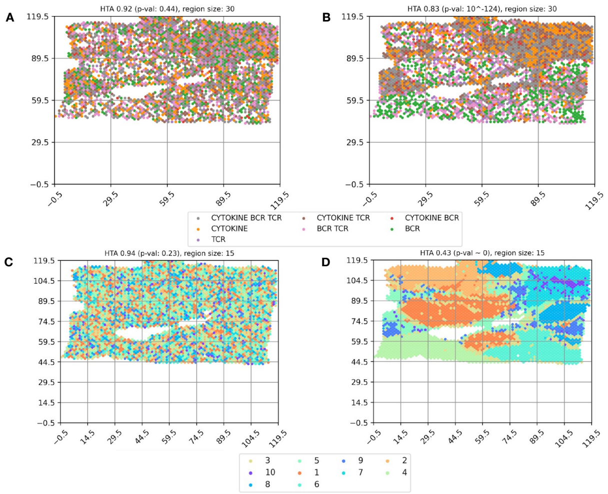


**Fig. 3.** Aggregating traits into meta-traits. In A-B, meta-traits are the following immune pathways: BIOCARTA_CYTOKINE_PATHWAY, BIOCARTA_TCR_PATHWAY, BIOCARTA_BCR_PATHWAY. Each pathway's enrichment values in each Visium spot was computed using the Bioconducters' GSVA package, and then binarized using the median threshold, as described in the main text for similar analyses. In C-D meta-traits are the cluster IDs obtained from 10X Genomics' Loupe Browser when selecting K-means with $k = 10$. (A), (C) represent a sample from the corresponding null models, while (B), (D) represent the actual data. The tissue is the same breast cancer tissue used in the main text.
